# A Multicenter Confirmatory Randomized-Controlled Study of rhNRGβ1 Protein Replacement Therapy in a Murine Model of NF2-related Schwannomatosis

**DOI:** 10.64898/2026.07.31.741963

**Authors:** Michael Reuter, Susann Groth, Johanna Schleep, Lars Björn Riecken, Lisa Schindler, M. Juliane Jung, Venkat Sundaram, Emilio Cirri, Nadine Pömpner, Lisa Wedekind, Julia Palm, André Scherag, Ruth Stassart, Robert Fledrich, Reinhard Bauer, Helen Morrison

## Abstract

**Background:** Previous exploratory studies identified recombinant human Neuregulin-1 β (rhNRGβ1) as a promising therapeutic strategy for inhibiting the growth of Nf2-deficient schwannomas by promoting cellular differentiation. Because robust confirmation across independent laboratories is essential for advancing promising preclinical findings toward clinical translation, we conducted a multicenter, randomized, controlled confirmatory study under stringent preclinical standards.

**Methods:** In a pre-registered trial (DOI: 10.17590/asr.0000304), 216 mice (Nf2-flox;P0-Cre;Nefh- Cre) were randomized at three independent research sites. Following a standardized sciatic nerve crush, mice received systemic rhNRGβ1 (10 µg/kg) or vehicle for 13 weeks. Rigorous quality measures included double-blinding, standardized surgery, centralized data management, and an automated Fiji macro for objective nerve thickness quantification (Primary Outcome). Secondary molecular outcomes included Western blot and in-depth, quantitative proteomics and phosphoproteomics. All methods were SOP-based for reproducible and comparable results across the three study centers

**Results:** The primary confirmatory analysis revealed no reduction in nerve thickness in the rhNRGβ1 group (p_best case imputation_ = 0.076 and p_worst case imputation_ = 0.533). Secondary analyses via quantitative Western blotting and DIA proteomics demonstrated that core biochemical markers of Schwann cell differentiation (MBP, ERBB2) remained unchanged across all centers. Based on the absence of macroscopic or primary biochemical effects, further histological analysis was omitted to avoid scientific redundancy. High-depth profiling of a predefined 60-protein functional marker panel confirmed a remarkably stable tumor proteome across all replication sites and both sexes, with no evidence of coordinated changes in key downstream oncogenic signaling pathways (Hippo/YAP, mTORC1, and RTK-Ras-MAPK) or metabolic signaling cascades. These findings indicate an absence of measurable target engagement under our tested dosing regimen, potentially reflecting pharmacokinetic or tissue-delivery limitations rather than an invalidation of the underlying biological pathway.

**Conclusion:** Despite high statistical power and rigorous methodology, this study could not confirm rhNRGβ1 as a robust therapeutic candidate for schwannoma growth arrest or shrinkage. These findings suggest that previously reported therapeutic effects were either highly context-dependent or could not be reproduced under adequately powered, rigorously controlled experimental conditions. As underpowered preclinical studies are more susceptible to random biological variation, our results highlight the importance of sufficient sample sizes alongside robust experimental design. Our study underscores the value of trial-like methodological standards in preclinical therapeutic evaluation to identify ineffective interventions (dead ends) early and strengthen translational decision-making. Although we could not confirm the previously reported efficacy of rhNRGβ1, the multicenter framework established here provides a methodological benchmark for robust preclinical testing in translational oncology, with the potential to improve reproducibility and the success of therapies progressing to early-phase clinical trials. From a translational perspective, these findings provide a robust foundation for optimizing future rhNRGβ1-based therapeutic approaches through improved dosing, delivery routes, and treatment schedules.

## Introduction

Neurofibromatosis type 2-related schwannomatosis (NF2-SWN) is a hereditary tumor syndrome characterized by the development of multiple benign, yet debilitating, Schwann cell-derived tumors along the peripheral and cranial nerves. These tumors, caused by bi-allelic loss of the NF2 tumor suppressor gene (encoding for Merlin), lead to progressive neurological deficits, including hearing loss and motor impairment [1], [2]. Currently, systemic pharmacological interventions are limited, often relying on repurposed anti-angiogenic agents with variable efficacy and significant side effects. Consequently, identifying novel, mechanistically grounded therapeutic targets remain a priority in NF2-SWN research [3], [4].

A promising therapeutic avenue involves the restoration of proper axon-glial signaling. Previous research has shown that NF2 deficiency in axons leads to a critical reduction in membrane-bound Neuregulin β1 (NRGβ1) type III, an essential instructive signal for Schwann cell differentiation. This deficit locks Schwann cells in a de-differentiated, proliferative state, creating a microenvironment conducive to schwannoma development. In a previous exploratory pilot study, we demonstrated that systemic and local administration of recombinant human NRGβ1 (rhNRGβ1) could functionally substitute for this loss, significantly reducing tumor growth and promoting Schwann cell maturation in mice [5].

Despite encouraging pilot findings, the field of translational research faces a significant "reproducibility crisis” [6]. Many promising preclinical candidates fail in clinical trials, often due to a lack of robustness in initial exploratory studies, which are frequently limited by small sample sizes, lack of blinding, or single-center biases [7], [8]. To overcome these limitations and provide a definitive assessment of rhNRGβ1 as a clinical candidate, we conducted a prospective, well-powered, multicenter confirmatory randomized-controlled study [9], [10].

The design of this trial (NRG1-Protein Replacement Therapy (NRG1-PRT)) adheres to the highest standards of preclinical rigor, faithfully emulating the design of human clinical randomized-controlled trials to ensure the maximum level of evidence [11]. The study was fully pre-registered (DOI: 10.17590/asr.0000304) prior to the commencement of experiments, defining all primary and secondary endpoints. To minimize experimental bias, we implemented a multicenter approach involving three independent research sites (FLI Jena, JUH Jena and ULMC Leipzig). Standardized Operating Procedures (SOPs) were strictly followed across all sites to improve reproducibility of surgical procedures, drug administration, measurements and data collection. Furthermore, we employed mandatory randomization and blinding for both the treatment phase and the outcome analysis. The study was investigator-blinded; the operator and all assisting personnel were unaware of treatment allocation. All study data were centrally managed via a secure REDCap database, ensuring data integrity and facilitating a robust Intention-to-Treat (ITT) analysis (Figure 1).

**Figure 1.**
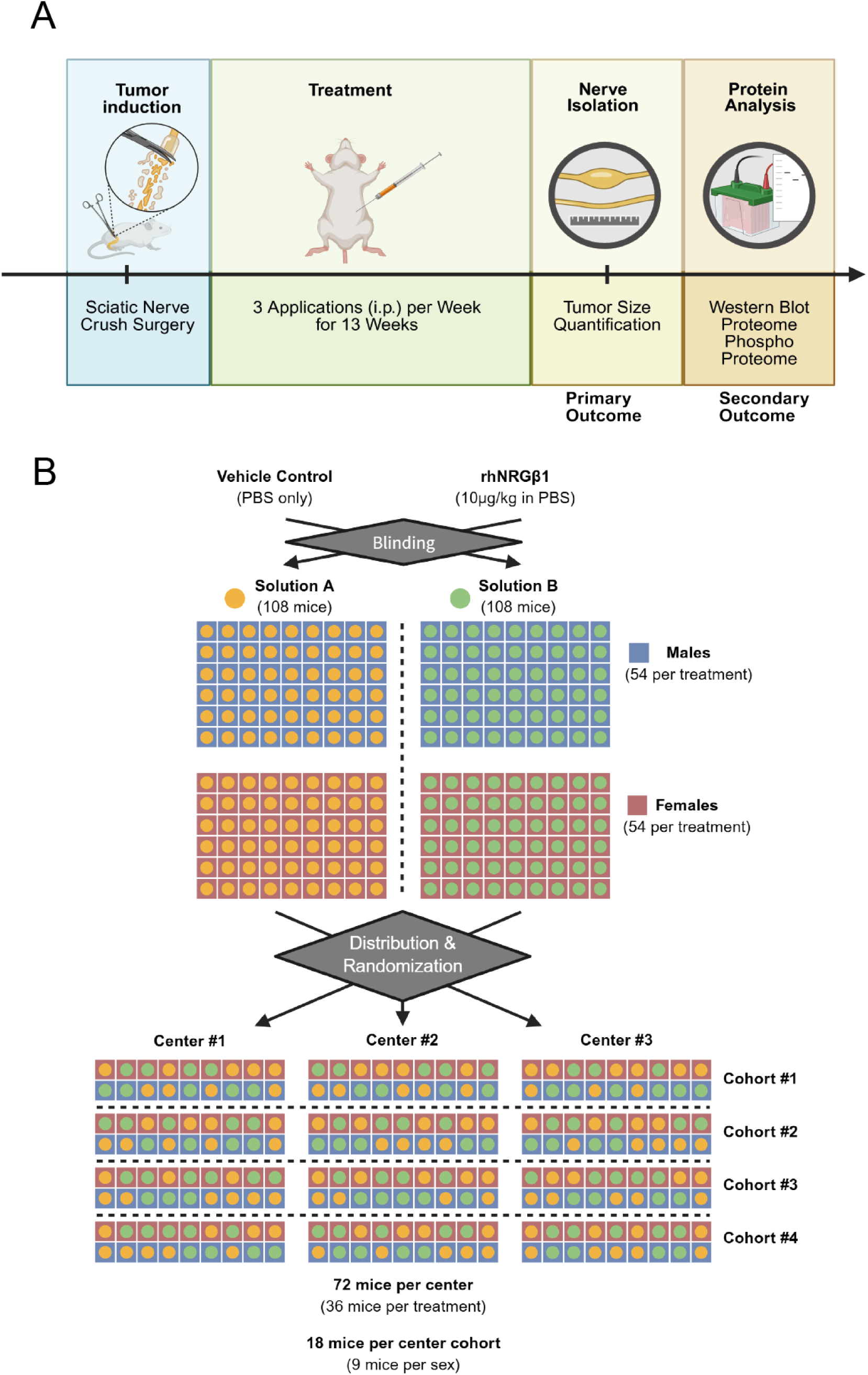
Experimental workflow and multicenter study design. **(A)** Following surgical tumor induction via standardized sciatic nerve crush, mice received intraperitoneal (i.p.) injections of either vehicle control or rhNRGβ1 three times per week for a total duration of 13 weeks. At the study endpoint, sciatic nerves were resected, photographed and evaluated using a semi-automated, image-based quantification macro to determine mean and maximum local nerve thickness (primary outcome). Lysates made from harvested whole nerves were split and processed for quantitative WB and high-depth DIA proteomics (secondary outcomes) **(B)** The randomized, double-blinded trial architecture comprised a total cohort of 216 mice, split evenly between the vehicle control (n = 108) and rhNRGβ1 (n = 108) treatment groups, with balanced stratification by sex (n = 54 males and n = 54 females per arm). Therapeutic solutions were fully blinded and coded as Solution A and Solution B to eliminate observation bias. The study population was stratified and distributed across three independent research centers, with each center evaluating a total of 72 mice divided into four distinct center cohorts of 18 mice each.

By employing a large-scale cohort (n = 216) and objective, semi-automated quantification methods, this study aims to validate the therapeutic potential of rhNRGβ1 in NF2-SWN. Beyond the primary morphological assessment of tumor growth, we utilized deep proteomics to evaluate the molecular bioactivity of rhNRGβ1 in vivo. Specifically, we leveraged a predefined 60-target proteomic panel representing critical signaling, metabolic, and cellular differentiation axes to comprehensively evaluate molecular target engagement and active tumorigenesis. This rigorous, confirmatory framework serves not only to test the preclinical efficacy of rhNRGβ1 but also to contribute to establishing a new standard for high-quality preclinical evaluation in order to improve the success rates in subsequent early clinical trials [12].

## Materials and Methods

### Study Design and Pre-registration

This study was designed as a prospective, multicenter, randomized, and double-blinded confirmatory trial. All experimental procedures and primary/secondary endpoints were defined a priori and formally pre-registered (DOI: 10.17590/asr.0000304). The study was conducted across three independent research sites in Germany: the Leibniz Institute on Aging – Fritz Lipmann Institute in Jena (referred to as “FLI”), the Jena University Hospital (referred to as “JUH”), and the Medical Experimental Center (MEZ) in Leipzig (referred to as “ULMC”). Data management, including randomization keys, animal welfare scoring, and primary outcome results, was documented centrally using a secure REDCap database to ensure data integrity and traceability.

### Animals and Ethics

Experiments were performed using a triple transgenic mouse line (Nf2-flox; P0-Cre; Nefh-Cre) on a mixed C57BL/6-FVB/N background. This genetic configuration results in the conditional deletion of the Nf2 gene in both Schwann cells and neurons of the peripheral nervous system, which enables tumor induction upon injury, accurately recapitulating the phenotypic hallmarks of human NF2-related schwannomatosis [3]. All experimental animals (fl/fl; tg/tg; tg/+) were centrally generated at an SPF barrier facility (FLI) and verified via PCR genotyping to ensure genetic consistency across all research sites. Mice of both sexes (12–16 weeks old, 16–45 g) were used and transferred to the experimental sites at least two weeks prior to surgery for acclimatization. Animal husbandry followed the 2012/63/EU guidelines, with mice housed in groups (2–5 per cage) under standardized conditions (12 h light/dark cycle) with *ad libitum* access to food and water. All procedures were approved by the respective governmental authorities (TLV Thuringia FLI-20- 012, UKJ-21-001 and Saxony TVV07-21 (ULMC)).

### Randomization and Blinding

To eliminate selection and observer bias, a rigorous blinding and randomization protocol was implemented.

1. **Randomization:** Animals were assigned to treatment groups using block randomization (block length of 18), stratified by sex and center, prepared by a biostatistician.
2. **Blinding of Solutions:** Recombinant human Neuregulin-β1 (rhNRGβ1; Reprokine #RKQ02297) and vehicle (PBS) were prepared and aliquoted by an independent individual not involved in surgery, injections, or data analysis. Solutions were coded as "Solution A" and "Solution B" with identical appearance. The blinding key was stored exclusively by the study monitor in a hidden REDCap form and was only unblinded after the completion of the primary outcome analysis (SOP NRG1-PRT-Solutions-EN).

### Standardized Sciatic Nerve Crush Injury

Surgical procedures were performed following a standardized SOP (NRG1-PRT-Surgery-EN). Under isoflurane anesthesia (2–3.5%), a unilateral nerve crush of the right sciatic nerve was performed, using hemostatic forceps (#13021-12; Fine Science Tools). The nerve was placed 1.4 mm from the tip, and a defined pressure was applied for 20 seconds by closing the forceps to the second interlocking click (referred to as “crush”). The left sciatic nerve remained intact as an internal control (referred to as “intact”).

### Therapeutic Intervention

Treatment commenced two days post-surgery (Wednesday) and continued for 13 weeks. Mice received intraperitoneal (i.p.) injections of either rhNRGβ1 (10 µg/kg) or vehicle (blinded as solution A and B) three times per week (Monday, Wednesday, Friday). Injection volumes were precisely calculated based on current body weight. Animal welfare was monitored three times per week using a multi-parameter scoring system (Figure 1).

### Primary Outcome: Nerve Diameter Quantification

Upon completion of the 13-week treatment (day of surgery + 92 days), mice were sacrificed and underwent the primary outcome analysis (n = 98–99 per treatment group). Both sciatic nerves were dissected and imaged on a black, non-reflective background alongside a size reference using a high-resolution single-lens reflex (SLR) camera (SOP NRG1-PRT-NerveSampling-EN).

Image analysis was performed in a fully automated, blinded batch process using a custom-made Fiji/ImageJ macro (NRG1-PRT-Macro-EN, Supplemental Data). The macro determined the local nerve thickness by measuring the diameter of the largest sphere that fits inside the nerve boundaries at every point along the nerve (Local Thickness algorithm). This objective method provided both mean and maximum nerve thickness values without researcher intervention. Importantly, no downstream processing, filtering, or manual adjustments were applied to the raw outputs generated by the macro.

To facilitate independent, blinded statistical evaluation, these raw, unaltered morphometric measurements were transferred directly to independent biostatisticians within a standardized master database (supplemental material). This dataset comprised animal identifiers, study center, biological covariates (sex, cohort), trial progression and attrition metrics, procedural success markers, the blinded group allocation, and the raw automated thickness parameters for both intact and crushed nerve segments (mean and maximum nerve thickness in mm).

### Secondary Outcomes: Biochemistry and Proteomics

According to the study design intact and crushed nerves from 94 mice (188 samples in total) were assigned to protein isolation for secondary outcome analyses, Western Blot and proteome analyses. While the protein yield was less critical for Western blot analyses, only those samples with a total yield of over 30 µg were used for further proteomic analyses (FLI: 54 mice, 26 females + 28 males; JUH: 66 mice, 32 females + 34 males; ULMC: 32 mice, 17 females + 15 males).

### Protein Extraction and Sample Preparation

Nerves were homogenized in TRIzol™ reagent (Thermo Fisher Scientific) according to the manufacturer’s instructions and a standardized protocol (SOP NRG1-PRT-TRIzol-EN). Following RNA extraction, proteins were precipitated from the organic phase with 100% ethanol. The resulting protein pellets were divided into two fractions: 70% for Western blot analysis and 30% for centralized proteomics. Pellets for Western blot were solubilized in a dedicated lysis buffer (containing SDS and DTT) and sonicated using a Bioruptor® (Diagenode; 10 cycles: 60 s ON / 30 s OFF). Samples were then heated at 95 °C for 5 minutes. Protein concentration was determined using a BCA protein assay to ensure equal loading.

### Western Blotting and Immunodetection

Western blot analysis was performed to quantify the expression of ERBB2 and Myelin Basic Protein (MBP) (SOP NRG1-PRT-WB-EN). Samples were separated by SDS-PAGE using 8% gels for ERBB2 and 15% gels for MBP. Proteins were transferred onto nitrocellulose membranes using center-specific established transfer systems. To account for loading variations, membranes were stained with Ponceau S for total protein normalization immediately after transfer.

Membranes were blocked and then incubated with the following primary antibodies overnight at 4 °C:

- Anti-ErbB2 (Cell Signaling Technology; #4290, Clone D8F12; Rabbit; Dilution 1:500)
- Anti-MBP (Sigma Aldrich; MAB384; Mouse; Dilution 1:500)

Following primary incubation and washing, membranes were incubated with appropriate horseradish peroxidase (HRP)-conjugated secondary antibodies. Immunoreactive bands were visualized using a digital chemiluminescence imager.

### Center-specific Western blotting protocol modifications

To optimize signal quality at the study center ULMC Leipzig, several protocol parameters were adjusted while maintaining the overall workflow defined by the SOP. Equal SDS concentrations were ensured across all samples before electrophoresis. For MBP detection, 15% SDS-PAGE gels were run at room temperature using an extended electrophoresis time (120 V for approximately 2 h), whereas ERBB2 samples were separated on 8% gels for approximately 3 h at 80 V. Instead of Ponceau S staining, total protein normalization was performed using the No- Stain™ Protein Labeling Reagent (Invitrogen, Thermo Fisher Scientific; Cat. No. A44449). Membranes were blocked in either 5% non-fat milk or 2% BSA, depending on the target protein. Primary antibody incubation conditions were modified, with MBP detected using anti-MBP 1:1000 overnight at 4 °C in 2% non-fat milk, and ERBB2 detected using anti-ERBB2 1:500 for 48 h at 4 °C in 2% BSA/TBS-T. Secondary antibodies were diluted 1:5000, using 2% non-fat milk/TBS-T for MBP and 0.5% BSA/TBS-T for ErbB2. Chemiluminescent signals were acquired using freshly prepared ECL substrate for each membrane, and quantitative analysis was performed using TotalLab software with normalization to the total protein signal.

### Quantitative Analysis

Quantification was performed using the software established at the respective study center (Image Lab at the centers FLI, JUH and TotalLab at ULMC Leipzig). For each sample, the HRP signal intensity of the target protein (specifically the 17 kDa and 21.5 kDa isoforms for MBP) was normalized to the respective total protein signal from the Ponceau S stain. To facilitate multicenter comparison while maintaining blinding, results were calculated as ratios of the mean signal of "Solution A" versus "Solution B" for each individual membrane before unblinding.

### Proteomics and Phosphoproteomics Sample Preparation and Protein Digestion

The 30% protein fraction obtained from the TRIzol™ extraction (SOP NRG1-PRT-TRIzol-EN) was used for deep proteomic and phosphoproteomic profiling. Protein pellets were solubilized in lysis buffer (5% SDS, 50 mM DTT, 100 mM HEPES, pH 8) and sonicated using a Bioruptor Plus (Diagenode) for 10 cycles (60 s ON / 30 s OFF). Following solubilization, samples were boiled at 95 °C for 5 minutes and alkylated with 15 mM iodoacetamide (IAA) for 30 minutes in the dark.

Samples were acidified with phosphoric acid (final concentration 2.5%), and seven times the sample volume of S-trap binding buffer was added (100 mM TEAB, 90% methanol). Samples were bound either on 96-well S-trap micro plate (Protifi) (for muscles) or on S-Trap micro columns (Protifi) and washed three times with binding buffer. Trypsin in 50 mM TEAB pH 8.5 was added to the samples (1 µg per sample) and incubated for 1 h at 47 °C. The samples were eluted in three steps with 50 mM TEAB pH 8.5, elution buffer 1 (0.2% formic acid in water) and elution buffer 2 (50% acetonitrile and 0.2% formic acid). The digests were then desalted with Waters Oasis® HLB µElution Plate 30 µm in the presence of a slow vacuum. In this process, the columns were conditioned with 3x100 µL solvent B (80% acetonitrile; 0.05% formic acid) and equilibrated with 3x 100 µL solvent A (0.05% formic acid in milliQ water). The samples were loaded, washed 3 times with 100 µL solvent A, and then eluted into PCR tubes with 50 µL solvent B. The eluates were dried using a speed vacuum centrifuge (Eppendorf Concentrator Plus, Eppendorf AG, Germany) and stored at −20 °C. Samples were then reconstituted in 100 ul MS Buffer (water with 5% FA) and then desalted.

### Phosphopeptide Enrichment

30 µg of peptides were enriched using Fe(III)-IMAC cartridges (Agilent) in an automated fashion using the standard protocol from the AssayMAP Bravo Platform (Agilent Technologies). In short, Fe(III)-IMAC cartridges were first primed with 100 µl of priming buffer (100% ACN, 0.1% TFA) and equilibrated with 50 μL of OASIS elution buffer. After loading the samples into the cartridge, the cartridges were washed with OASIS elution buffer, while the syringes were washed with priming buffer. The phosphopeptides were eluted with 25 μL of 1% ammonia directly into 25 μL of 10% FA. The flowtrough was collected to measure non phosphorylated peptides. Samples were dried down with a speed vacuum centrifuge and stored at −20 °C until LC-MS analysis.

### LC-MS/MS Data Acquisition

For whole cell analysis, 10 µL of samples were transferred to a 96-well plate (Eppendorf twin.tec PCR LoBind skirted, Eppendorf AG, Germany) separated using a Vanquish Neo system (Thermo Scientific) equipped with a trap column (PepMap™ Neo Trap Cartridge, 5 mm length, 300 μm ID, 5 μm particle size) and an Ionopticks Aurora Rapid 8×75 (8 cm and 75 μg, 1.7 μm C18 particle size, IonOpticks) column heated at 55 °C. Loading of the trap was done in a combined control mode (flow 50 μl/min, 800 bar) with an automatic loading volume. A 15.5 minutes gradient (70SPD) was used, ramping from 4% to 8% B (ACN with 1% FA) in 0.5 minutes, then up to 28% in 9 minutes, to 42% B in 3.5 minutes, followed by a zebra washing (4 cycles, combined control) and a fast equilibration (flow 0.5 μl/min, 1400 bar, equilibration factor 4). The flow was kept at 300 nL/min during separation.The LC system was coupled to an Orbitrap Astral (Thermo Fisher Scientific) via a Proxeon nanospray source heated at 300 °C, and a spray voltage of 1.8 kV was applied. The radio frequency ion funnel was set to 40%.

Data were acquired in Data-Independent Acquisition (DIA) mode. For whole cell analysis, full MS1 scans (380–980 m/z) were acquired at a resolution of 240,000 FWHM. The default charge state was set to 2+. The filling time was set at maximum of 3 ms with limitation of 5 × 10^6^ ions (ACG 500%). DIA MS2 scans were acquired in the Astral Analyzer with 300 dynamic window width across the MS1 mass range, a scan range from an *m*/*z* of 150 to 2000 was chosen. Higher collisional dissociation fragmentation was 25%, the precursor accumulation time was set to 3 ms, the AGC target was set to 5 × 10^6^ ions (500%). Data were acquired with a time loop control of 0.6 seconds. For data acquisition and processing of the raw data Xcalibur 4.7 (Thermo) and Tune version 2.1 were used.

For phosphoproteomics analysis, peptides were separated using a Vanquish Neo system (Thermo Scientific) equipped with a trap column (PepMap™ Neo Trap Cartridge, 5 mm length, 300 μm ID, 5 μm particle size) and an Ionopticks Aurora Rapid 8×75 (8 cm and 75 μg, 1.7 μm C18 particle size) column heated at 55 °C. Loading of the trap was done in a combined control mode (flow 50 μl/min, 800 bar) with an automatic loading volume. A 13 minutes gradient (100SPD) was used, ramping from 4% to 8% B (ACN with 1% FA) in 0.2 minutes, then up to 40% in 10.4 minutes, to 50% B in 0.2 minutes and finally to 99% B in 0.5 minutes and kept for 1.7 minutes, followed by a zebra washing (4 cycles, combined control) and a fast equilibration (flow 0.5 μl/min, 1400 bar, equilibration factor 4). The flow was kept at 300 nL/min during separation. The LC system was coupled to an Orbitrap Astral (Thermo Fisher Scientific) via a Proxeon nanospray source heated at 300 °C, and a spray voltage of 1.8 kV was applied. The radio frequency ion funnel was set to 40%. DIA parameters were kept the same as for whole cell analysis except for the variable windows, that were adapted based on phosphopeptides distribution.

### Data Processing and Bioinformatic Analysis

DIA raw data were analyzed using the directDIA pipeline in Spectronaut v.20 (Biognosys, Switzerland) with BGS settings besides the following parameters: Protein LFQ method= QUANT 2.0, Proteotypicity Filter = Only protein group specific, Major Group Quantity = Median peptide quantity, Minor Group Quantity = Median precursor quantity, Data Filtering = Qvalue, Normalizing strategy = Local Normalization. The data were searched against a species specific (*Mus musculus*, 16,747 entries, v. 210401) and a contaminants (247 entries) Swissprot database. The identifications were filtered to satisfy FDR of 1% on peptide and protein level. The data were then exported and further analyzed in RStudio using MSStats [13] removing sparse precursors and with a cutoff for missing values of 0.5. The protein quantification and differential expression table used for volcano plots generation and Principal Component Analysis (PCA), respectively, using R version 4.1.3 and RStudio server version 1.1.463. Protein groups were considered as significantly enriched if they displayed a q value < 0.01 and average log2 ratio > 0.58.

For phosphoproteomics analysis, the data were searched with the following modifications: Carbamidomethyl (C) (Fixed) and Oxidation (M), Acetyl (Protein N-term), Phospho (STY) (Variable). PTM localization probability was set to 0.75 and consolidation of phosphosites was sum based. A maximum of 2 missed cleavages for trypsin and 5 variable modifications were allowed. The identifications were filtered to satisfy FDR of 1% on peptide and protein level. Relative quantification at phosphosite level was performed in Spectronaut for each paired comparison using the replicate samples from each condition. The data (candidate table) and data reports (protein quantities) were then exported and further data analyses and visualization were performed with RStudio using in-house pipelines and scripts. To select significant phosphosites, a log2FC cutoff of 0.58 and a q value < 0.05 were defined. For identifying the most regulated kinases and the corresponding regulated top phosphosites, a modified version of PhosR [14] was used.

## Statistical Analyses

Sample size of this trial was calculated (n = 216) to provide 90% power to detect a biologically relevant difference (Cohen’s d = 0.5) between the treatment groups based on prior pilot data at a significance level α = 0.05 (two-sided). The primary confirmatory analysis was performed on the modified ITT analysis set (see below) using the Brunner-Munzel test [15], which addresses the Behrens-Fisher problem in non-parametric rank-based distributions. Note that the estimated relative effect in the non-parametric setting denotes the probability that a randomly chosen measured primary outcome from a mouse in one group exceeds a randomly chosen outcome in the control group. This yields an effect size on a 0–1 scale where 0.5 indicates no difference and values away from 0.5 indicate a directional shift. We report the estimated relative effect with its 95% confidence interval (CI) in addition to the corresponding (two-sided) p-values. Missing values of the primary outcome were handled via worst-case/best-case imputation (necessary for 9/207 animals – see Figure 2) and we applied a significance level α = 0.05 (two-sided). Mixed-effect models were used as sensitivity analyses to account for center-specific random effects and sex as a covariate. All analyses were conducted using R version 4.1.3 and RStudio server version 1.1.463.

**Figure 2.**
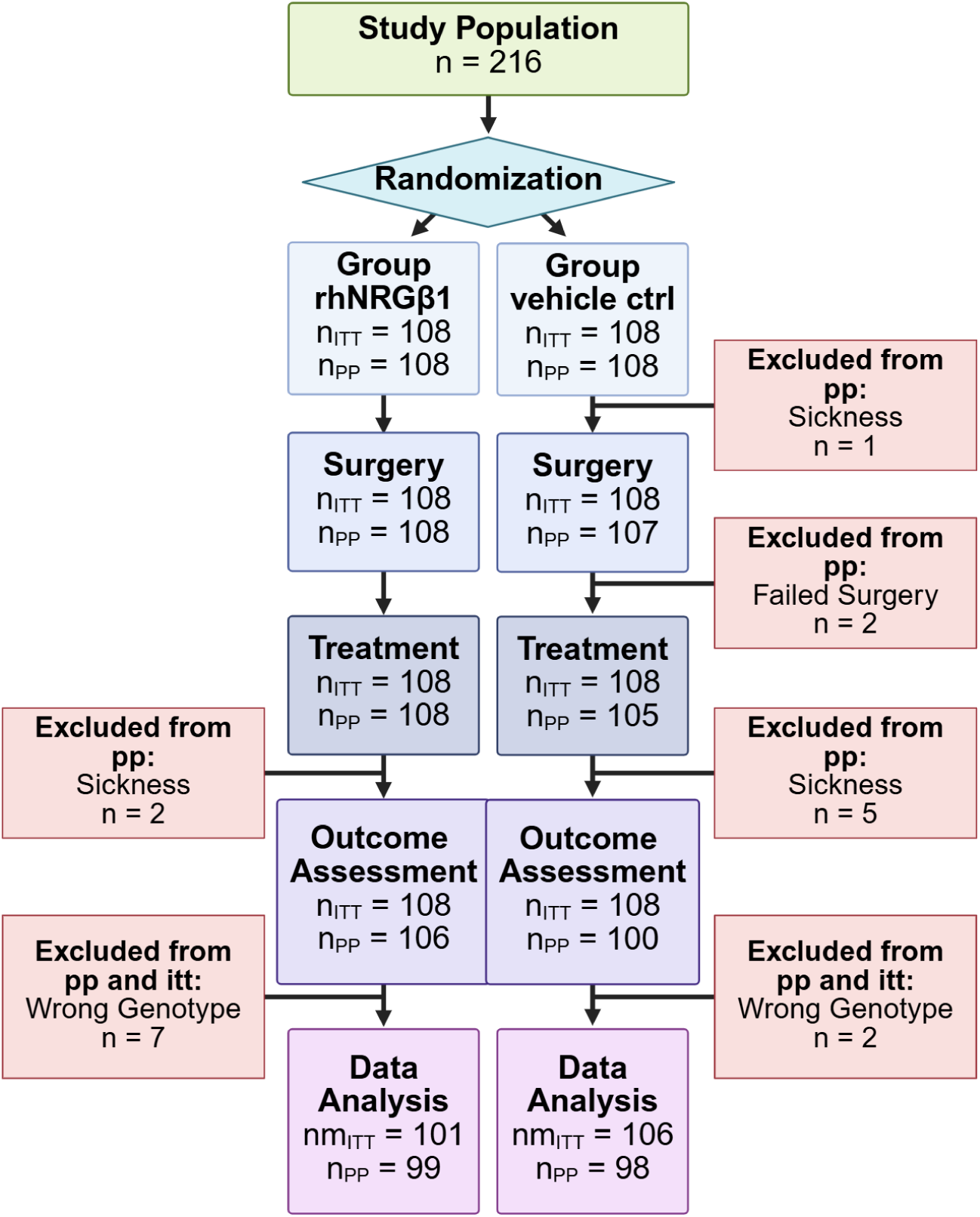
Experimental flowchart detailing (modified) Intention-to-Treat (ITT) and Per-Protocol (PP) population. Schematic representation of the multicenter study design from initial randomization to final analysis. The ITT population includes all randomized mice that entered the treatment phase, serving as the primary baseline to preserve statistical power and randomization integrity across centers. The modified ITT (nm ITT) population includes all randomized animals but specifically excludes those identified with incorrect genotypes. The Per-Protocol (PP) population restricts the final analysis strictly to individual animals that completed the full 13-week study period without experiencing premature health exclusions, surgical dropouts, or technical data collection failures. The specific numbers of excluded animals and the biological reasons for attrition are provided at each trial tier.

## Results

### Study Attrition and Data Integrity

To ensure the highest level of data integrity, we monitored experimental adherence across all three centers via a centralized REDCap database. Of the 216 mice initially enrolled, we defined a modified ITT (nm ITT) analysis set comprising 207 mice, which excluded 9 animals subsequently identified with incorrect genotypes. Although all mice underwent initial genotyping prior to randomization, our quality management protocol required post-sampling re-genotyping to verify animal identities. Importantly, this genotype verification was completed prior to any treatment unblinding, preventing potential selection bias. The final Per-Protocol (PP) population analysis set addressed additional exclusions for protocol deviations, including technical failures and early dropouts, resulting in a subgroup of 197 mice. Overall, the attrition rate was minimal and balanced across treatment groups and centers; a detailed summary is provided in Figure 2.

### Primary Outcome: Confirmatory Analysis of Nerve Thickness

The primary objective was to determine if systemic rhNRGβ1 treatment reduces the mean thickness of the schwannoma-bearing nerves. We employed the Brunner-Munzel test to account for potential variance heterogeneity in rank-based distributions across the multicenter cohort and used worst-case/best-case imputation to handle missing values.

In the primary confirmatory analysis (modified ITT analysis set; n = 207), rhNRGβ1 treatment did not lead to a reduction in mean nerve thickness compared to the vehicle group. The estimated relative effect was 0.57 (95% CI: 0.49 0.65; p = 0.076) under best-case imputations. This finding was consistent across two different imputation methods for missing values (Figure 3 and Table 1).

**Figure 3.**
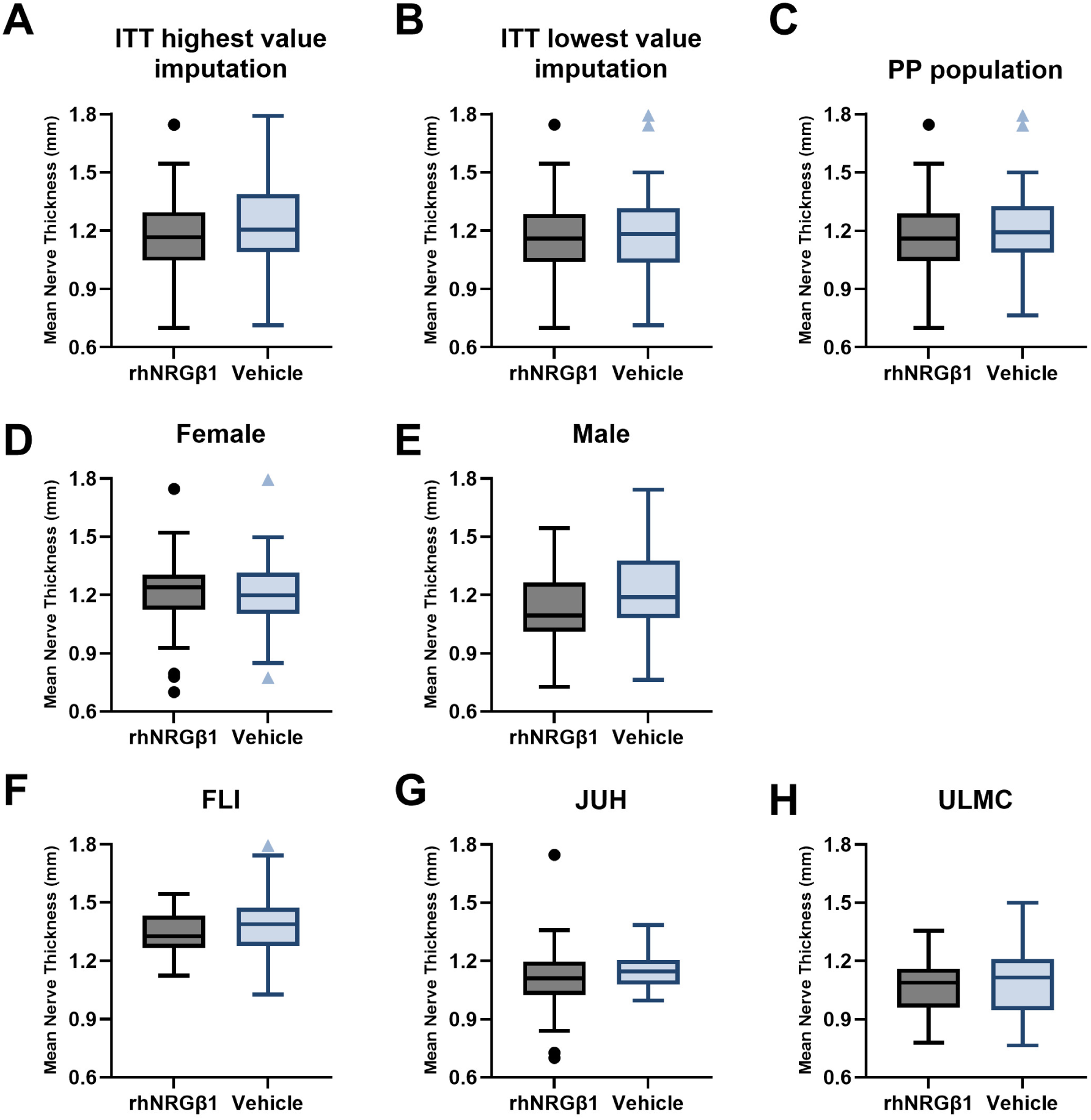
Semi-automated morphometric quantification of tumor-bearing sciatic nerve thickness across all centers. Box plots display mean local nerve thickness (mm) evaluated via an automated investigator-independent Fiji macro. **(A and B)** ITT population analyzed using highest-value **(A)** or lowest-value **(B)** imputation methods (nm_ITT_ = 101–106 per group). **(C)** Per-Protocol (PP) population (n_PP_ = 98–99 per group). **(D and E)** PP population stratified into female and male cohorts (n = 49–50). **(F, G, and H)** Exploratory breakdowns split by individual study centers: JUH, FLI, and ULMC (n = 64–67 per center).

**Table 1.** Results of the Brunner-Munzel test comparing treatment groups.

| Type of analysis | Population | n | Effect estimator [95% CI] | p value |
| --- | --- | --- | --- | --- |
| Confirmatory | ITT highest value imputation | 207 | 0.57 [0.49; 0.65] | 0.0761 |
| Confirmatory | ITT lowest value imputation | 207 | 0.53 [0.45; 0.60] | 0.5325 |
| Exploratory | PP | 197 | 0.56 [0.48; 0.64] | 0.1674 |

The exploratory analysis in the PP population further confirmed this result (n = 197, effect estimator 0.56 (95% CI: 0.48. 0.64; p = 0.167), indicating that the lack of efficacy was not driven by outliers or protocol deviations (Figure 3C). Similarly, secondary morphological parameters, such as maximal nerve thickness and normalized thickness (subtraction of mean or maximal thickness of the intact nerve from the crushed nerve), revealed no evidence for differences between groups (Table 2).

**Table 2.**
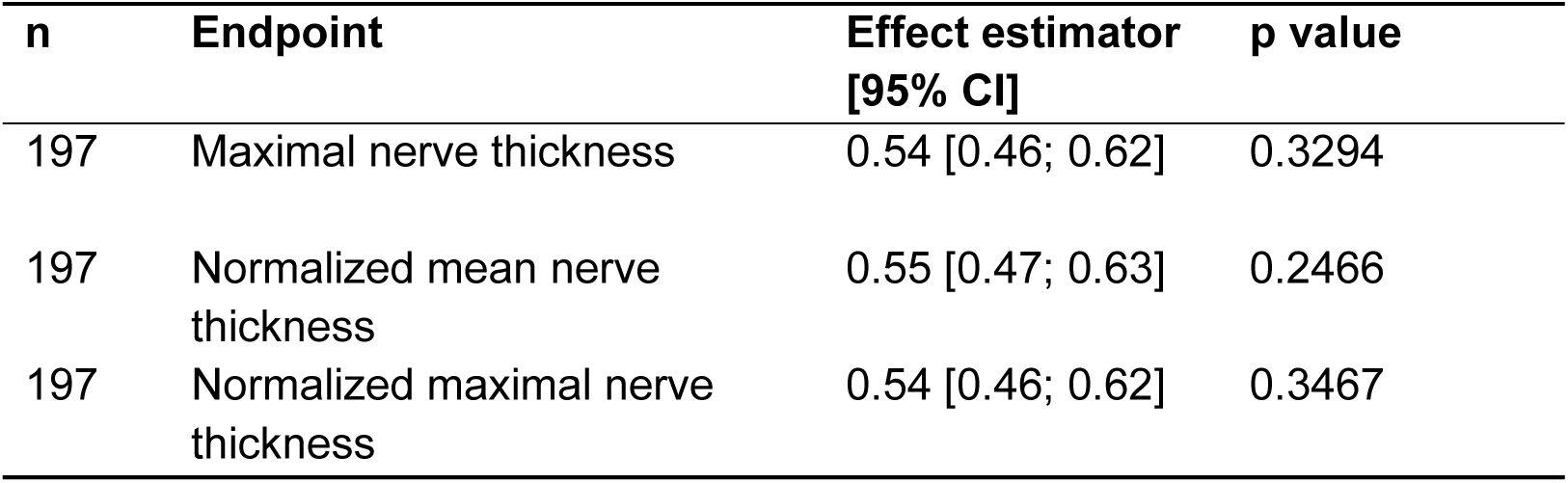
Results of the exploratory analysis comparing treatment groups.

| n | Endpoint | Effect estimator [95% CI] | p value |
| --- | --- | --- | --- |
| 197 | Maximal nerve thickness | 0.54 [0.46; 0.62] | 0.3294 |
| 197 | Normalized mean nerve thickness | 0.55 [0.47; 0.63] | 0.2466 |
| 197 | Normalized maximal nerve thickness | 0.54 [0.46; 0.62] | 0.3467 |

### Exploratory Subgroup and Center-Specific Analysis

To investigate potential sources of variance, we performed subgroup analyses by sex or research site again using non-parametric tests (Figure 3D–H and Table 3). These explorations revealed that there might be an effect on reduced mean nerve thickness under systemic rhNRGβ1 relative to vehicle in male mice (effect estimator 0.63; p = 0.025), particularly male mice at the JUH site (effect estimator 0.72; p = 0.020). However, this finding was not supported by a parametric linear mixed-effects model accounting for research center as a random effect (p = 0.189 for the treatment effect in the PP analysis set). Furthermore, inclusion of a treatment-by-sex interaction also provided no evidence that the treatment effect differed between males and females (p = 0.080), suggesting that the subgroup finding should be interpreted with caution.

**Table 3.** Statistical Discrepancy Analysis: Local Rank-Based Subgroup Trends vs. Global Hierarchical Linear Mixed-Effects Model for Sciatic Nerve Thickness.

| Analysis Type | Population / Stratum | Statistical Test | Main Statistical Output | p value |
| --- | --- | --- | --- | --- |
| Subgroup | Males (All Centers Combined) | Brunner-Munzel | Effect Estimator: **0.63** [95% CI: 0.52; 0.74] | **0.0246** |
| Site-Specific Subgroup | Males at **JUH** Center | Brunner-Munzel | Effect Estimator: **0.72** [95% CI: 0.54; 0.90] | **0.0202** |
| Site-Specific Subgroup | Males at **FLI** Center | Brunner-Munzel | Effect Estimator: **0.56** [95% CI: 0.35; 0.77] | 0.5469 |
| Site-Specific Subgroup | Males at **ULMC** Center | Brunner-Munzel | Effect Estimator: **0.63** [95% CI: 0.42; 0.84] | 0.2252 |
| **Global Pre-Registered Model** | **All Subjects** (PP Population) | **Linear Mixed-Effects Model** (Fixed: Treatment + Sex; Random: Center) | Treatment B Estimate: 0.028 (SE: 0.021) | **0.1885** |

### Molecular Assessment of Target Engagement and Bioactivity (Secondary Read-outs)

While the primary "macroscopic" outcome – the quantification of nerve thickness – revealed no evidence for a reduction in tumor growth, the secondary outcomes were designed to evaluate whether rhNRGβ1 effectively engaged its molecular targets or if the biological failure occurred at a downstream level. Based on previous exploratory findings [5], our working hypothesis was that rhNRGβ1 would act as a "differentiation switch", triggering proliferating repair SC to redifferentiate [16] into myelinating SC, leading to a significant upregulation of Myelin Basic Protein (MBP) and a concomitant normalization (downregulation) of the ERBB2 receptor tyrosine kinase.

### Western Blot Analysis of Key Differentiation Markers

To test our pre-registered hypothesis that rhNRGβ1 would act as a molecular "differentiation switch" we performed quantitative Western blotting for the primary target molecules defined in our protocol: ERBB2 and MBP. We selected these specific markers to monitor the molecular remodeling of the nerve tissue; ERBB2 functions as the obligate co-receptor mediating NRG1 signaling in the Schwann cell lineage, whereas MBP serves as a canonical structural marker of mature, myelinating glia. If rhNRGβ1 successfully induced differentiation within the dedifferentiated, proliferating tumor environment, we expected to see a robust upregulation of myelin structural proteins – particularly the isoforms associated with active myelination – accompanied by a significant down-regulation or stabilization of ERBB2 expression due to receptor trafficking or differentiation-induced feedback loops.

Western blot analysis across all participating centers and cohorts showed no significant differences between the treatment groups. Both the 17 kDa and the 21.5 kDa isoforms of MBP – the latter being a marker for active re-myelination – remained at levels comparable to vehicle-treated controls. Similarly, the expression of ERBB2 showed no significant reduction (Figure 4A representative blots). To achieve reliable inter-blot comparison, we initially normalized target band intensities to total protein loading via Ponceau S (or No-Stain™ Protein Labeling Reagent for ULMC samples), and subsequently to the mean value of internal, vehicle-treated controls on each respective blot (4 blots per center with a total of n = 47). These molecular data confirm that the primary pre-registered molecular targets of Schwann cell maturation were not globally affected, mirroring the negative macroscopic results (Figure 4B).

**Figure 4.**
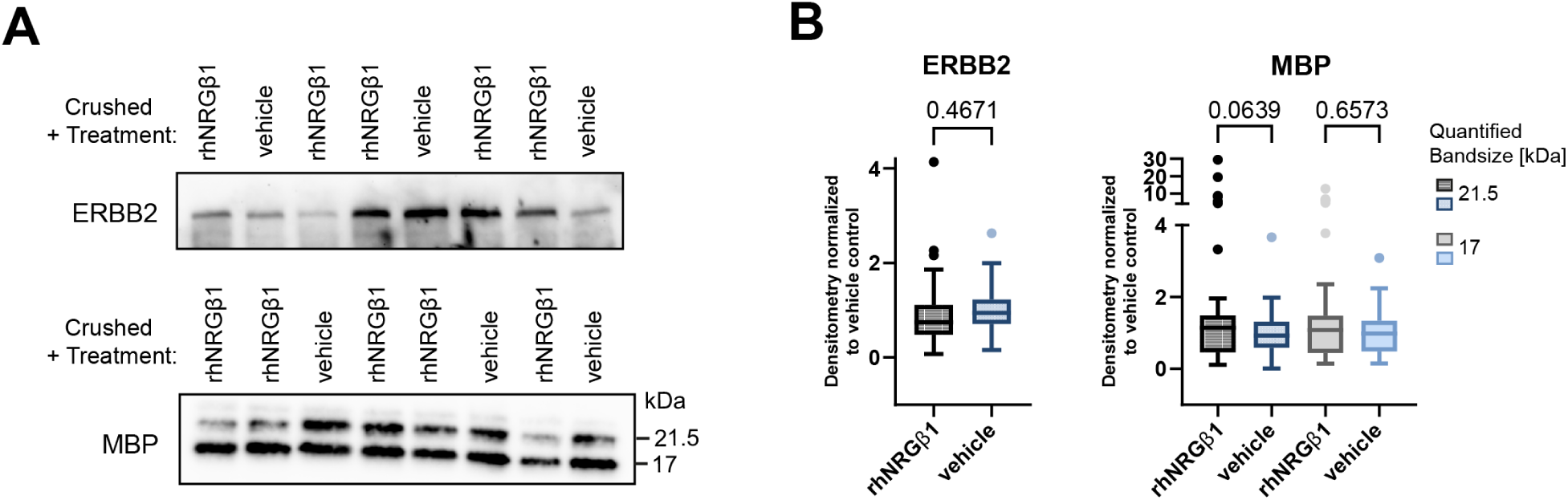
Quantitative Western blot analyses of ERBB2 and Myelin Basic Protein (MBP) in tumor-bearing nerves. **(A)** Representative immunoblots for ERBB2 (upper panel), and MBP (lower panel) made with nerve lysates from JUH cohort 4. **(B)** Relative quantification of ERBB2 (left panel) and MBP (right panel) across all study centers and cohorts, normalized to total protein loading (Ponceau S) and intra-blot vehicle control means to display center- and cohort-specific quantitative distributions (n = 47).

This biochemical finding was further cross-validated by the high-depth proteomic data (Figure 5A). Both MBP and ERBB2 were robustly detected within the global nerve proteome; however, their expression levels showed no significant differential regulation (log2FC) when comparing rhNRGβ1-treated nerves to the PBS vehicle control. This lack of primary target modulation remained consistent across all cohorts, completely independent of sex or nerve status (crushed or intact).

**Figure 5.**
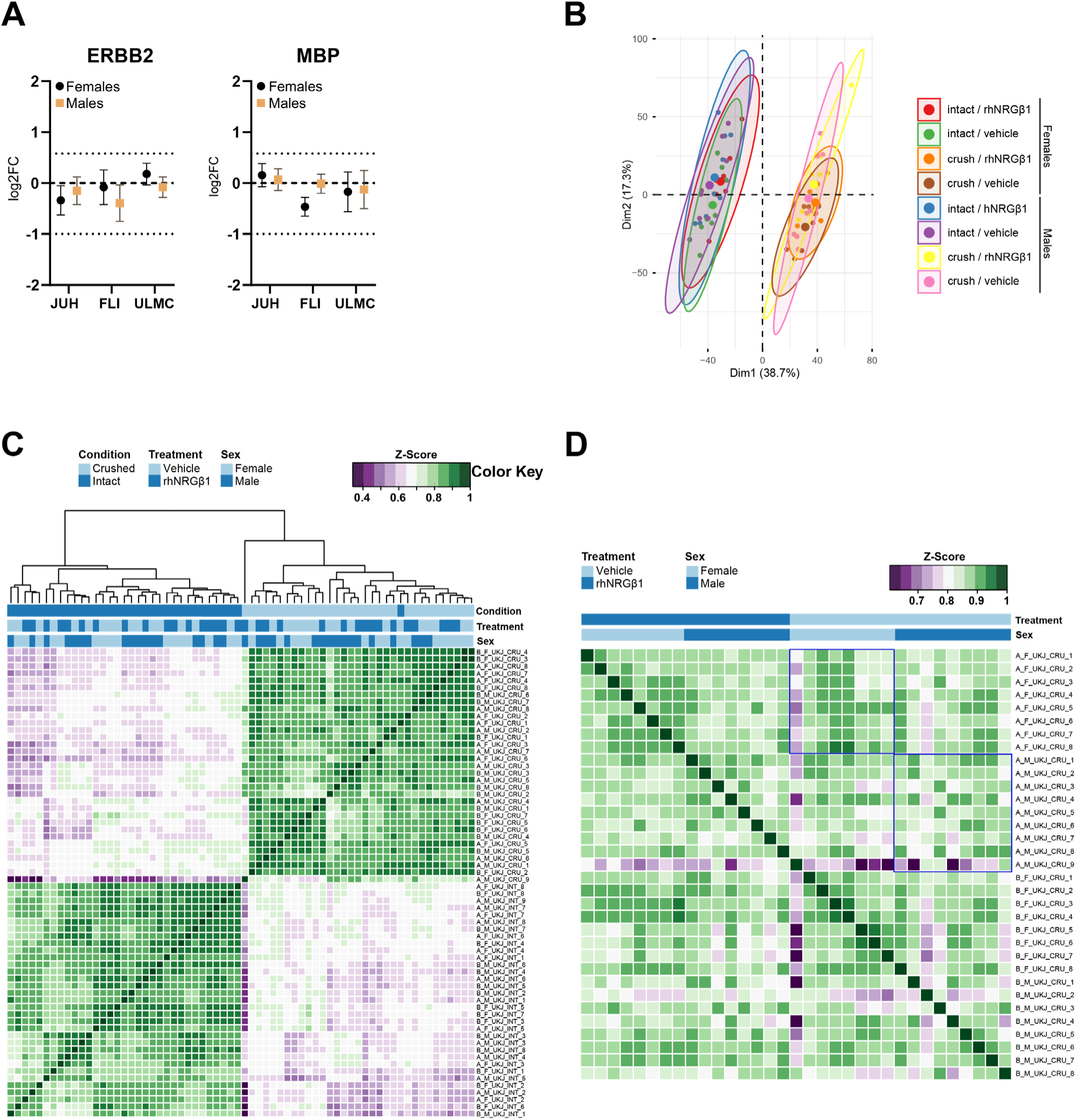
Quantitative DIA proteomic profiling, sample dimensionality, and cohort correlation analyses. **(A)** log2 fold changes (log2FC) of the pre-registered marker proteins ErbB2 and MBP across male and female cohorts, stratified by the three participating study centers. **(B)** Principal Component Analysis (PCA) of global proteomic profiles from the JUH center, segregated by nerve condition, treatment group, and sex. **(C)** Pearson correlation matrix demonstrating global statistical associations across all processed JUH samples. **(D)** Pearson correlation matrix exclusively for tumor-bearing sciatic nerve samples of the JUH cohort. The relevant comparison groups (female vehicle vs rhNRGβ1 and male vehicle vs rhNRGβ1 are framed in blue.

### Proteomic Profiling and Bioactivity Verification

To further explore potential changes at a deeper molecular level, we conducted high-depth DIA-proteomics. Principal Component Analysis (PCA) demonstrated that the global proteomic landscape of rhNRGβ1 treated nerves was almost identical to that of vehicle controls, with overlapping clusters for both sexes and with the nerve crush condition being the main driver of differences (Figure 5B). This is also reflected by the sample correlation matrix, where samples with a strong correlation are indicated in green, while less strong correlation is shown in purple (Figure 5C). Based on this the nerve tumor (crush) condition is the main parameter driving differences between crushed vs intact samples. Thus, sex and treatment (rhNRGβ1, vehicle) showed up as strong correlation. Showing that neither sex nor treatment led to major differences among the samples. Also, the Pearson Correlation used here, revealed two major clusters containing crushed vs intact nerve samples, while there was neither a rhNRGβ1 treatment nor a sex specific clustering, again illustrating strong similarity among these samples.

When focusing only on the crushed (tumor carrying) nerves in a correlation matrix, we again do not see differences among rhNRGβ1 vs vehicle treated nerves (Figure 5D), illustrating again the strong similarity among these samples. This molecular uniformity was further corroborated by untargeted phospho-proteomic profiling, which revealed an identical absence of differential phosphorylation states or treatment-induced signaling shifts (Data provided in supplemental material). These findings directly mirror the total proteome landscape, confirming that rhNRGβ1 administration failed to trigger downstream kinase activation cascades.

Although the unbiased global proteome lacked widespread statistical significance across cohorts, we performed a targeted, hypothesis-driven evaluation of 60 predefined markers (Table 4). This allowed us to specifically interrogate key molecular axes driving nerve-relevant processes and active tumorigenesis, regardless of global threshold limitations.

**Table 4.** Functional classification and proteomic coverage of the preselected 60-marker panel.

| Category | Detected Targets | Non-Detected Targets | Biological Reflection / Relevance | Ref |
| --- | --- | --- | --- | --- |
| **Cell Proliferation & Apoptosis** | CASP3 | MKI67, BAX, BCL2 | Balances active mitotic division driving autonomous tumor growth against programmed cell death signaling activated during tissue damage. | [25], [26] |
| **Fibrosis and ECM Remodeling** | S100A4, COL1A1, FN1 | ACTA2 | Represents the injury-induced wound healing response and tissue scarring alongside the formation of a supportive desmoplastic stroma. | [27] |
| **Hippo/YAP-Signaling** | YAP1 | WWTR1, CYR61, CTGF | Tracks the primary proximal mechanotransduction pathway hyperactivated by NF2 loss, which acts as the major transcriptional engine preventing Schwann cells from exiting the cell cycle during regeneration. | [28] |
| **Immune Infiltration & TME** | CD68, MRC1, CD163, CD38, CD200, SLC7A2, ITGB2, VCAM1 | S100A9 | Maps leukocyte and macrophage recruitment, blending the acute inflammatory cleanup of myelin debris with the establishment of an immunosuppressive tumor microenvironment. | [29], [30] |
| **mTORC1 Pathway & Metabolism** | SIRT2, MTOR, RPS6KB1, RPS6 | None | Reflects the translational machinery activation and metabolic rewiring required to meet the high energetic demands of both axonal/glia repair and volumetric schwannoma expansion. | [31] |
| **Receptor Tyrosine Kinases (RTKs) & Signaling** | ERBB2, ERBB3, STAT3, STAT6, EGFR, PDGFRA, PDGFRB, NGFR, SHH, AXL, IGF1R | ERBB4, NRG2, NRG1, JUN, VEGF, MST1R | Monitors the aberrant membrane accumulation of growth factor receptors and autocrine/paracrine signaling loops caused by the loss of Merlin-mediated contact inhibition in NF2-related schwannomatosis | [32] |
| **Schwann Cell Differentiation & Neuronal Markers** | MBP, MPZ, S100B, GFAP, APOE, APOD, FLRT3, NCAM1, L1CAM, NEFH | None | Evaluates the phenotypic transition between mature myelinating glia, dedifferentiated repair Schwann cells clearing debris, and the permanently blocked progenitor state of tumorigenesis. | [16] |
| **Tumor Suppressors, ERM Complex & Cellular Architecture** | NF2, PRKCQ, EZR, RDX, MSN, CD44, SMARCB1 | LZTR1 | Establishes the structural tissue baseline and submembrane actin scaffolding, mapping the mechanical collapse of cell-to-cell adhesion and contact inhibition that initiates schwannomatosis. | [33], [34] |

Out of these 60 predefined markers, 45 targets were found to be expressed and reliably quantified in the proteome across our experimental groups, whereas the remaining 15 proteins fell below the analytical limit of detection (Figure 6).

**Figure 6.**
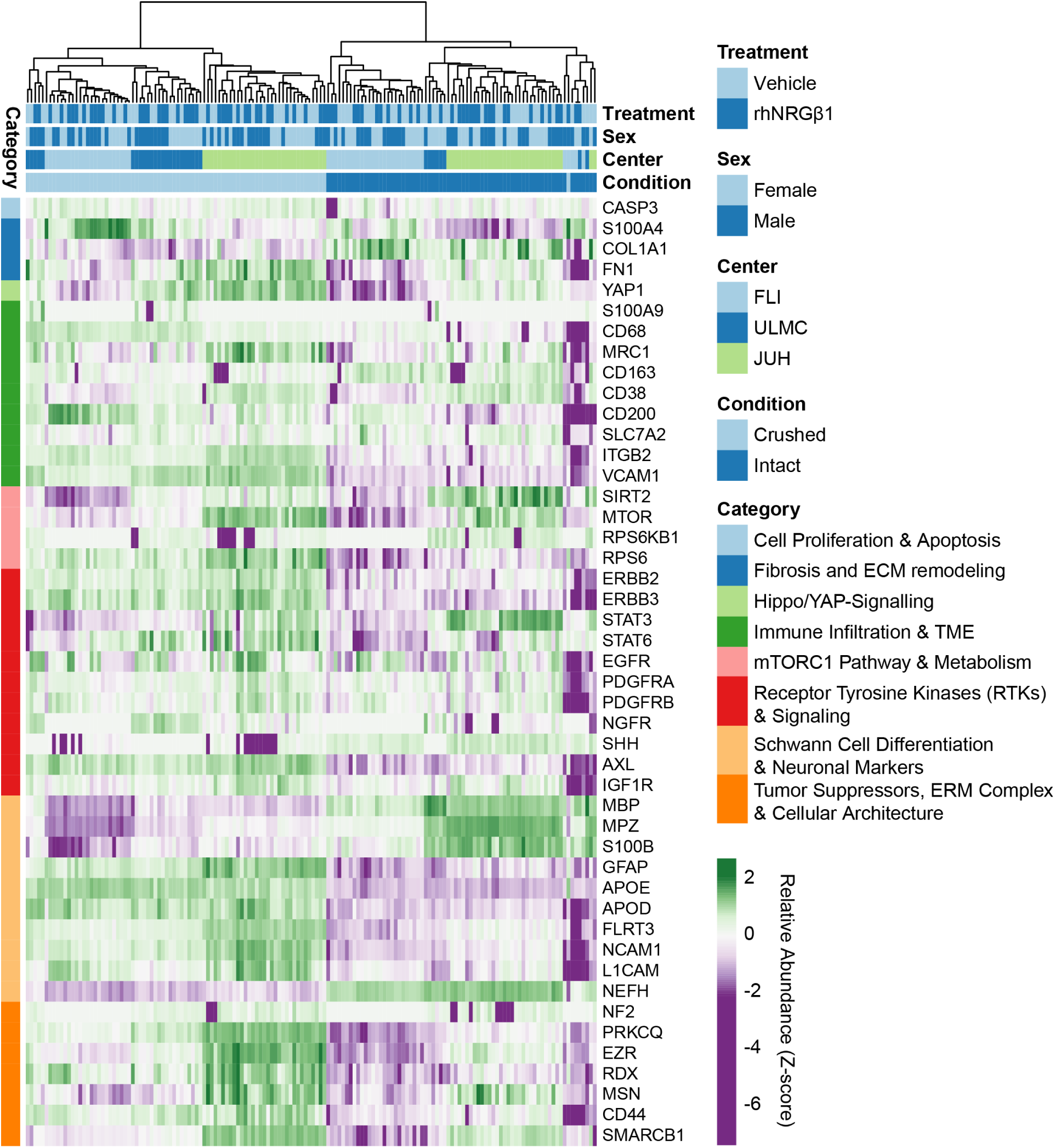
Heatmap and Pearson correlation analysis of a predefined 60-protein functional marker panel evaluating study-relevant biological pathways. Protein abundance levels across key signaling, metabolic, and cellular differentiation pathways relevant to nerve function are clustered using Pearson correlation and displayed independent of statistical significance. Expression profiles encompass all processed samples across all study centers, both sexes, and all nerve conditions (crushed/tumor-bearing and intact).

### Quantitative Proteomic Evaluation of Therapeutic Efficacy Across Study Centers

A central additional objective of this multicenter study design was to establish reproducible, harmonized proteomic readouts of therapeutic intervention across the three participating research sites (JUH, ULMC and FLI). Theoretically, identical experimental cohorts should yield convergent proteomic trajectories under therapeutic pressure. However, global differential analysis revealed notable divergence between the study centers, coupled with a highly limited overall drug effect. After applying stringent multiple-testing corrections, the vast majority of predefined target proteins failed to achieve the statistical significance threshold (q < 0.05). This generalized lack of robust target modulation suggests that the intervention did not induce widespread, cross-center anti-tumor response within the injury-induced tumor or the surrounding nerve tissue.

### Heterogeneous Responses in Tumor-Bearing Nerves

When analyzing the tumor-bearing sciatic nerves (vehicle vs. rhNRGβ1), the proteomic outcomes bifurcated sharply by study center and sex:

### JUH Cohorts

At the JUH center, therapeutic intervention left the tumor proteome almost entirely unaltered. No single target approached multiple-testing significance, with all adjusted values remaining uniformly elevated (q > 0.60). The only notable trend was a weak, nominal upregulation of the metabolic sensor SIRT2 exclusively in female tumor-bearing nerves (log2FC = 0.26, p = 0.0127, q = 0.629). Downstream translational elements (MTOR, RPS6KB1, RPS6) and the core structural component of the SWI/SNF chromatin-remodeling complex (SMARCB1) remained completely static.

### FLI Cohorts

In contrast to the quiet baseline observed at JUH, the female tumor-bearing cohort at the FLI site exhibited a localized, statistically robust tissue remodeling signature. Most notably, the integral myelin structural protein MPZ was significantly downregulated upon treatment (log2FC = −0.52, p = 8.46 10^-6^, q = 0.019), representing one of the few true FDR-significant changes in the study. This structural demyelination profile was supported by corresponding nominal drops in MBP (log2FC = −0.47, p = 0.016, q > 0.9) and the axonal control NEFH (log2FC = −0.59, p = 0.005, q = 0.67), alongside a nominal accumulation of the unmyelinated glial marker S100B (log2FC = 1.12, p = 0.029, q > 0.9). Conversely, male tumor-bearing mice at the FLI site showed no such structural alterations, displaying only minor nominal shifts in EGFR (log2FC = −1.34, p = 0.011, q = 0.87) and FN1 (log2FC = 0.36, p = 0.038, q > 0.9).

### ULMC Cohorts

Aligning closely with the highly stable molecular landscape observed at the JUH site, the ULMC center demonstrated a predominantly uniform proteomic profile between treatment groups across both sex cohorts. The 60-protein panel verified that downstream oncogenic signaling pathways, such as the Hippo/YAP axis (YAP1) and the metabolic MTORC1 cascade (MTOR, RPS6KB1, RPS6), stayed firmly at baseline. Widespread coordinated pathway modulation was absent, with the dataset yielding only isolated, cohort-specific fluctuations, such as a nominal shift in the immune microenvironment marker CD163 within the second cohort (log2FC = −0.59, p = 0.280, q = 0.049).

In summary, the statistical reality of the dataset indicates that the therapeutic intervention did not achieve a uniform, reproducible therapeutic effect. The presence of opposing, center-specific signatures – such as the structural myelin alterations isolated exclusively to the FLI tumor cohorts contrasted against the remarkably static molecular baselines maintained across both the JUH and ULMC platforms – powerfully illustrates how localized technical or environmental variances can eclipse subtle drug effects, heavily penalizing the global q-values across all evaluation sites.

### Justification for the Omission of Histological Analysis

The original study protocol (pre-registration DOI: 10.17590/asr.0000304) envisioned a comprehensive histopathological analysis part, including immunohistochemical quantification of S100b, MPZ, P75, and KI67. However, following the unequivocally negative findings of both the primary macroscopic endpoint and the secondary biochemical and proteomic screens, the steering committee decided to omit the histological evaluation.

## Discussion

This study represents, to our knowledge, the first well-powered, multicenter preclinical confirmatory randomized controlled trial explicitly designed to evaluate the therapeutic potential of rhNRGβ1 protein replacement therapy in a mouse model of NF2-related schwannomatosis. Moving far beyond the single-center exploratory framework that dominates preclinical research, we established a standardized pipeline across three completely independent research institutions (FLI Jena, JUH Jena, and ULMC Leipzig). Our primary macroscopic endpoint based on the semi- automated quantification of nerve thickness and our secondary molecular readouts (ERBB2 and MBP expression) collectively demonstrated that the systemic administration of 10 µg/kg rhNRGβ1 three times weekly did not achieve an arrest of injury-induced schwannoma growth.

While these results diverge from the observations of therapeutic efficacy in our initial pilot study [5], this outcome should not be interpreted as a biological failure of the rhNRGβ1-based replacement strategy. Instead, this trial provides an invaluable diagnostic calibration point, clarifying the strict pharmacokinetic limits, dosing sensitivities, and delivery parameters required to successfully advance this candidate forward.

## Paradigmatic Rigor: Establishing Clinical-Grade Trial Design in Preclinical Models

A major strength of this study is the implementation of a clinical trial-inspired preclinical framework, introducing methodological standards long established in randomized human clinical trials to basic mouse model-driven schwannomatosis research. The reproducibility crisis in translational research, particularly in neuroscience, has highlighted how promising therapeutic candidates often fail because early findings are generated in single-center studies with limited sample sizes and methodological limitations that increase the risk of bias and false-positive results. To address these challenges, the NRG1-PRT study was deliberately designed to emulate the rigor of late-phase randomized controlled trials [12].

Key features, including multicenter execution, prospective protocol registration, centralized double-blinded randomization, predefined endpoints, adequate statistical power, and standardized operating procedures, were incorporated to maximize internal validity, minimize experimental bias, and ensure reproducibility. As a key component of this framework, the complete study protocol was prospectively preregistered before study initiation (DOI: 10.17590/asr.0000304), thereby limiting opportunities for selective outcome reporting and post hoc statistical inflation (p-hacking). Collectively, our findings demonstrate that rigorously designed preclinical studies can yield informative negative results that are equally valuable for translational decision-making. By reliably identifying ineffective therapeutic strategies, such frameworks help direct resources toward more promising candidates while establishing a methodological benchmark for robust, transparent, and reproducible preclinical evaluation in translational oncology.

## Reconciling the Discrepancy: Sample Size and the "Winner’s Curse"

When interpreted within the context of a rigorous confirmatory trial, the discrepancy between our current neutral findings and the therapeutic efficacy observed in our initial exploratory study can be understood constructively. The original pilot study demonstrated that systemic and local rhNRGβ1 administration could promote Schwann cell differentiation and reduce schwannoma growth. However, these exploratory cohorts included relatively small sample sizes (n = 5–12 animals per group), making them inherently more susceptible to random sampling variation and the “Winner’s Curse”, whereby early positive studies tend to overestimate treatment effects because only the most favorable results achieve statistical significance and are preferentially reported [7], [17], [18], [19].

In contrast to the pilot study, which was susceptible to the “Winner’s Curse”, the present confirmatory study was specifically designed to address these limitations as a preregistered, adequately powered, multicenter trial involving more than 200 mice. Increasing sample size reduces not only random variation but also the influence of biological and technical variability inherent to in vivo studies – including inter-animal differences in disease progression and treatment response, environmental factors, and measurement variability – allowing genuine treatment effects to be distinguished more reliably from background experimental noise. While larger cohorts cannot compensate for systematic bias, preregistration, randomization, double blinding, and standardized multicenter procedures further minimize bias and selective outcome reporting. Together, these methodological safeguards substantially reduce the likelihood that observed effects reflect chance findings rather than true biological activity, thereby mitigating the “Winner’s Curse”.

Importantly, expanding the study across multiple centers also captures the biological and environmental heterogeneity that more closely reflects real-world experimental and, ultimately, clinical settings. Within this framework, the global hierarchical linear mixed-effects model did not confirm an overall treatment effect, despite isolated subgroup trends observed in local analyses. These findings suggest that although rhNRGβ1 may retain context-dependent biological activity, the current systemic delivery paradigm is not sufficiently robust to consistently suppress schwannoma growth across a heterogeneous multicenter population.

## Preserving the Translational Viability of rhNRGβ1: Pharmacokinetic Refinement and Delivery Optimization

Crucially, these neutral results do not invalidate the rhNRGβ1 replacement hypothesis, as the fundamental axon-glial signaling axis underlying Nf2 pathobiology remains biologically sound. In NF2-related schwannomatosis, the loss of Merlin downregulates axonal Neuregulin-1 type III, trapping surrounding Schwann cells in a permanently dedifferentiated, proliferative loop [20], [21]. Because our multicenter biochemical and proteomic profiles demonstrated that the core differentiation markers Myelin Basic Protein (MBP) and ERBB2 remained static, this neutral outcome points directly to a pharmacokinetic or delivery ceiling rather than an inherent failure of the biological target. To transition rhNRGβ1 into an effective tumor-suppressive therapy, future validation phases must optimize key delivery parameters, beginning with a transition from intraperitoneal (i.p.) injections to direct intravenous (i.v.) administration. While the current i.p. route subjects the recombinant protein to variable peritoneal absorption kinetics and first-pass clearance, direct i.v. bolus injections can maximize plasma peak concentrations to more efficiently exploit receptor-mediated active transport mechanisms across the blood-nerve barrier [22].

## Tiered Evaluation and Scientific Pragmatism

Finally, our adherence to a strict, data-driven "stop-signal" for the histopathological analysis part underscores the clinical maturity of our preclinical framework. Given that both the automated nerve diameter calculations and the high-depth global proteomic screens confirmed that the primary differentiation machinery remained unaltered, proceeding with large-scale, descriptive immunohistochemical analysis for S100β, MPZ, or P75 was deemed scientifically unjustified.

By avoiding the "sunk cost fallacy" [23], this pragmatic pivot allows us to responsibly conserve critical mouse tissue biobanks, institutional resources, and research funding. These saved assets can be directly reallocated into executing the optimized intravenous, high-dose, and continuous-infusion delivery schedules required to fully realize the translational potential of rhNRGβ1 replacement therapy.

## Conclusion and Implications for NF2 Research

In conclusion, although the primary therapeutic hypothesis was not supported under the present experimental conditions, this study establishes a rigorous methodological framework for future preclinical NF2 research. The systematic integration of centralized data capture and management, prospective protocol preregistration, centralized randomization, double-blinding, standardized multicenter procedures, and investigator-independent automated image quantification using Fiji minimized potential sources of bias and strengthened confidence in the robustness of the negative findings. The study design and statistical analysis were further supported by independent biostatisticians, and the project benefited from scientific exchange with the DECIDE project at the Berlin Institute of Health QUEST Center [24], which promotes best practices for confirmatory preclinical research beyond the specific questions addressed here.

From a translational perspective, these findings provide an evidence-based foundation for decision-making by indicating that the current rhNRGβ1 delivery strategy does not warrant clinical translation. By preventing the advancement of ineffective therapeutic approaches, rigorous preclinical evaluation helps safeguard patients while ensuring that limited research funding and clinical resources are directed toward more promising interventions. Rather than closing the chapter on rhNRGβ1 replacement, the present study provides a reliable reference point for future investigations aimed at optimizing dosing, delivery routes – including direct intravenous administration – and treatment schedules to fully evaluate its differentiation-inducing potential.

## Supporting information

Primary Outcome Data Table

## Author Contributions

Conceptualization/Study design: HM, LBR, MR, AS; Biostatistical analyses (primary outcome): AS, LW, JP; Study Monitor: LBR; Study preregistration & SOP writing: LBR, MR, MJJ, SG; Design, setup and steward of REDCap project: SG (with input from MR, LBR, MJJ for design); Research Data Management: LBR, MR, LS, VS; Proteome analyses pipelines with subsequent data processing and bioinformatic analysis: EC, NP; Conduction of experiments: MR, VS, MJJ, SG; Data analysis and curation (secondary outcomes): MR, SG, JS, VS, LS, LBR; Writing original manuscript: MR, JS, SG, HM, EC, AS, JP; Reviewing & editing: all authors; Figures: MR, SG, JS (Figs. 1–6 + supplementary); LBR (design Fig.1); AS, JP (design Fig. 2); Funding/Grant writing: HM, LBR, AS, RS, RB; Animal license writing: MR, MJJ, RB, RF; Supervision: HM, RS, RF, RB.

## Acknowledgments

The authors thank the DECIDE project at the QUEST Center for Responsible Research, Berlin Institute of Health, Charité-Universitätsmedizin Berlin, Berlin, Germany for their support during the project, the FLI Core Facility Proteomics, the dedicated staff at the FLI Mouse Facility, the FLI Core Facility Life Science Computing (Fabian Monheim, REDCap administration), the FLI Core Facility Technology Transfer (Sonja Schätzlein); and all involved facilities at the JUH and ULMC for their essential support and infrastructure. We are also extremely grateful to Michelle Grünewald, Uta Papke, Marina Zahid, Rose Zimmer, and Alexander Gloria for their outstanding technical assistance. We also thank Anita Barzegar-Fallah, Georgia Daraki and Luisa Ricciardi for their support during the injection program. Figures 1 and 2 created in BioRender. Reuter, M. (2026 Fig. 1A: https://BioRender.com/swrw5pp, Fig. 1B: https://BioRender.com/5dc8eg9, Fig. 2: https://BioRender.com/k160dq1

## Data and materials availability

All SOPs referenced in the manuscript are attached to the pre-registered study “NRG1-PRT - rhNRGß1 protein replacement therapy for treatment of Schwann cell nerve sheath tumors”, DOI 10.17590/asr.0000304 and will be publicly available soon after the manuscript’s publication.

All MS data, source code and raw data will be archived in general-purpose repositories with free access following their peer review and upon publication.

