## Supplementary Data for "A Multicenter Confirmatory Randomized-Controlled Study of rhNRGβ1 Protein Replacement Therapy in a Murine Model of NF2-related Schwannomatosis"

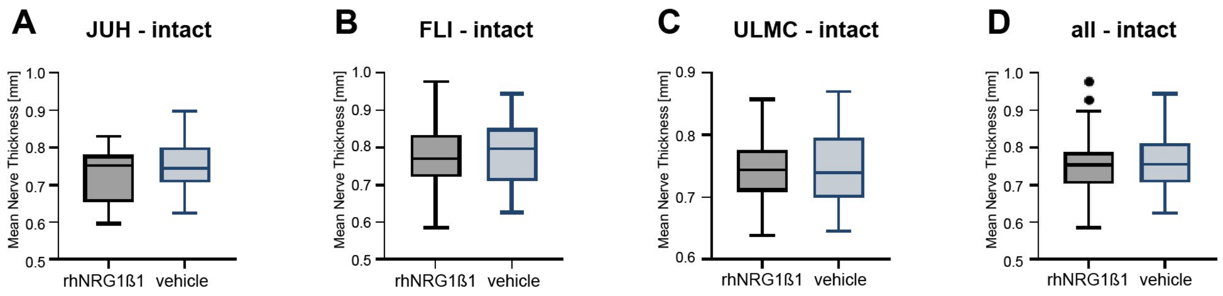


**Supplementary Figure 1. Semi-automated morphometric quantification of intact sciatic nerve thickness.** Box plots display mean local nerve thickness (mm) evaluated via an automated, investigator-independent Fiji macro. **(A–C)** Intact nerve thickness profiles analyzed by individual study centers: JUH (A), FLI (B), and ULMC (C) (n = 64–67 per center). **(D)** Merged analysis combining data across all three centers to display the global quantitative distribution.

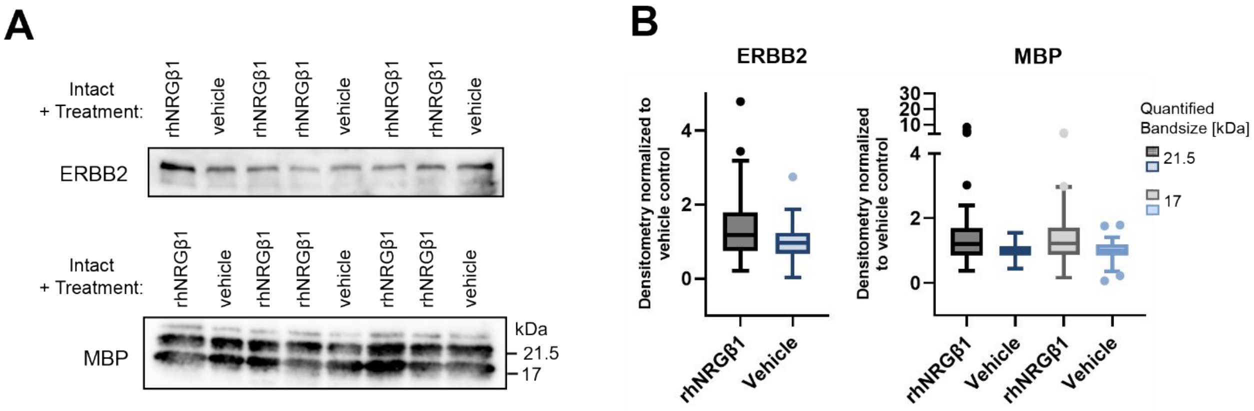


**Supplementary Figure 2. Quantitative Western blot analyses of ERBB2 and Myelin Basic Protein (MBP) in intact nerves. (A)** Representative immunoblots for ERBB2 (upper panel) and MBP (lower panel) made with nerve lysates from JUH cohort 4. **(B)** Relative quantification of ERBB2 (left panel) and MBP (right panel) across all study centers and cohorts, normalized to total protein loading (Ponceau S) and intra-blot vehicle control means to display center- and cohort-specific quantitative distributions (n = 47)

### FIJI-Macro for Nerve thickness analysis (IJM-Format):

```
// Recursively opens files and measures nerve thickness

requires("1.32f");
run("Line Width...", "line=3");
run("Colors...", "foreground=white background=black selection=yellow");
run("Clear Results");
roiManager("reset");
 close("*");

//Selection of Folder with images for analysis
dir = getDirectory("Choose Source Directory ");
dir2 = dir + "/Analysis/";
File.makeDirectory(dir2);
dir_res = dir + "/Results/";
File.makeDirectory(dir_res);

//Scaling - only done once for all images made in a row
path = File.openDialog("Select an image for scaling.");
 open(path);
setTool("line");
waitForUser("Please mark 1cm and press Ok");
while (!is("line")) {
      waitForUser("Error, please draw a line and mark 1cm!");
}
run("Set Scale...", "known=10 unit=mm global");

 close("*");

//Batch processing of all images in the selected folder
count = 1;
prefix = "mask-";
listFiles(dir);

function listFiles(dir) {
    list = getFileList(dir);
    for (i=0; i<list.length; i++) {
        if (endsWith(list[i], "/"))
            listFiles(""+dir+list[i]);
        else {
            print((count++) + ": " + dir + list[i]);
            open(dir + list[i]);

            c = 1;
            while (nImages > 0)  {
                //analysis starts here

orgImageName = getTitle();

//Dialogbox for user input of mouse ID
  title = "ID";
    Dialog.create("New Image");
  Dialog.addString("Animal-ID:", title);
  Dialog.show();
  title = Dialog.getString();
k=0;
do{       //loop for measuring both nerves

//Selection of nerve
setTool("rectangle");
waitForUser("Please, select the nerve you want to measure.");
while (selectionType() != 0)                         {   //make sure we have got a
rectangular selection
      waitForUser("Error, please draw a rectangle!");  }

//Dialog to select if nerve is crushed or intact
  treatment = newArray("Crush", "Intact");
  Dialog.create("Treatment");
  Dialog.addRadioButtonGroup("Treatment:", treatment, 1, 2, "Intact");
  Dialog.show();
```

```
  treat = Dialog.getRadioButton();

//pre-processing 1. cut out 2. pre-process
run("Duplicate...", "title=" + title);
run("Scale Bar...", "width=5 height=12 font=62 color=White background=None
location=[Lower Left] bold overlay");
run("Duplicate...", "title=" + title + "-" + treat);
run("Enhance Contrast...", "saturated=0.01 normalize");
run("8-bit");
run("Enhance Contrast...", "saturated=0.01 normalize");
run("Median...", "radius=20");
run("Duplicate...", " ");
run("Auto Threshold", "method=RenyiEntropy white");
run("Options...", "iterations=1 count=1 black do=Nothing");
run("EDM Binary Operations", "iterations=8 operation=open");

setTool("line");
waitForUser("Please check mask and press Ok. \n If corrections are necessary, draw a
line and press delete to adjust selection. "); //if selection was not perfect, user is
able to adjust this
run("Adjustable Watershed", "tolerance=30"); //the tolerance might be adjusted
depending on the image resolution

run("Analyze Particles...", "size=20000-Infinity pixel show=Masks exclude add"); //the
size might be adjusted depending on the image resolution. This is to exclude all
smaller pieces and to focus only the nerve.
run("Grays");
run("Local Thickness (masked, calibrated, silent)");
run("Set Measurements...", "area mean standard median min max display redirect=None
decimal=3");
roiManager("Measure");

//save as PNG

                saveAs("PNG", dir2 + title + "_" + treat + "-" + list[i] + "-" + c +
".png");
                 selectWindow(title);
                 roiManager("Select", 0);
                 run("Draw", "slice");
                 saveAs("PNG", dir2 + title + "_" + treat + "-" + list[i] + "-mask-" +
c + ".png");
                 roiManager("Delete");
                selectWindow("Results");
                saveAs("Measurements", dir_res + "Results.csv"); //save as Results
after every image

            //repeat for 2nd nerve.
      selectWindow(orgImageName);
          //reply = getBoolean("Have both nerves been analyzed?", "Yes", "No");
    k=k+1;
} while(k<2)
 close("*");
                c++;
            }
        }
    }

    //selectWindow("Results");
    //saveAs("Results", dir_res + "Results_final.csv");
}
```
